# Epoxyeicosatrienoic acids and sEH inhibition prevent cardiac dysfunction in CVB3-induced myocarditis by positively regulating type I interferon signaling

**DOI:** 10.1101/2023.02.03.527086

**Authors:** Zhou Zhou, Min Zhang, Chengcheng Zhao, Xu Gao, Zheng Wen, Junfang Wu, Chen Chen, Jiong Hu, Ingrid Fleming, Dao Wen Wang

## Abstract

Myocarditis is a challenging inflammatory disease of the heart and better understanding of its etiology is needed to develop specific drug therapies. Epoxyeicosatrienoic acids (EETs), active molecules synthesized by cytochrome P450 enzymes from arachidonic acids and hydrolyzed to less active dihydroxyeicosatrienoic acids by soluble epoxide hydrolase (sEH), have been attributed anti-inflammatory activity. Here, we investigated whether EETs have immunomodulatory activity and exert protective effects on coxsackie B3 virus (CVB3)-induced myocarditis. Viral infection altered eicosanoid epoxide and diol levels in both patients with myocarditis and, in the murine heart, and correlated with the increased expression and activity of the sEH after CVB3 infection. Administration of a sEH inhibitor prevented CVB3-induced cardiac dysfunction and inflammatory infiltration. Importantly, EET/sEH inhibitor treatment attenuated vial infection/ improved viral resistance by activating type I interferon (IFN) signaling. At the molecular level, EETs enhanced the interaction between glycogen synthase kinase 3β (GSK3β) and TANK-binding kinase 1 (TBK1) to promote IFN-β production. Our findings revealed that EETs and sEH inhibitors prevent the progress of CVB3-induced myocarditis, particularly by promoting viral resistance by increasing IFN production.

## Introduction

Myocarditis is clinically and pathologically defined as a heart-specific inflammatory disease caused by a range of infections and non-infectious triggers [1, 2]. Myocarditis is regarded as the third leading cause of cardiovascular death in young athletes [3, 4] and its most common cause is viral infection e.g., by enteroviruses, parvovirus B19, and adenoviruses [5–7]. Indeed, up to 40% of dilated cardiomyopathy is associated with viral infection or inflammation in the myocardium [1, 8]. Viral myocarditis is characterized by three phases: an acute phase during which viruses infect and replicate in the body, a subacute phase with a marked immune response and a chronic phase of dilated cardiomyopathy [6, 7, 9]. Encouragingly, improvement in the treatment of fulminant myocarditis (FM); a condition with severely compromised hemodynamics and a high mortality rate [10], have helped to decrease mortality rates from over 50% to less than 5% in distinct clinical centers [10].

Murine myocarditis induced by the coxsackie B3 virus (CVB3) is an excellent model for dissecting inflammatory processes in the heart, i.e., the direct cytopathic effects of the virus as well as the subsequent host immune response [7, 11, 12]. While adverse immune responses can contribute to tissue injury, persistent viral proliferation directly affects cardiomyocyte viability and contributes to the progression of myocarditis [1, 11, 13]. At the moment, there is no approved antiviral therapy for patients with viral myocarditis [5, 7, 13]. Given that viral infections lead to the production of interferons (IFNs); which impact adaptive immune responses and are important for host defense against viruses [14, 15], viral clearance with IFN-β has been linked to the maintenance of cardiac function in animal models as well as in patients with myocarditis [16–21].

Epoxyeicosatrienoic acids (EETs), are bioactive metabolites of arachidonic acid (AA) that are generated by cytochrome P450 (CYP) enzymes, that have been attributed anti-hypertensive, anti-inflammatory and cardioprotective actions [22–24]. Endogenous EETs are rapidly metabolized to less inactive diols or dihydroxyeicosatrienoic acids (DHETs) by the soluble epoxide hydrolase (sEH) (encoded by *EPHX2*), and the use of sEH inhibitors is an effective way of stabilizing endogenous EET levels and enhancing their beneficial effects [25, 26]. A role for EETs as potent anti-inflammatory mediators has been suggested by a series of studies in different models of inflammation. Mounting evidence indicates that EETs inhibit the expression of inflammatory molecules by reducing the activation of NF-κB [27–30], and elevated levels of EETs (in particular 14,15-EET) can suppress the cardiac inflammatory response by preventing macrophage activation [31]. However, the role, if any, of sEH inhibition in myocarditis is not known. Therefore, in this study we determined whether sEH inhibition and EETs play a role in myocarditis and whether inhibiting the sEH to increase EET levels can prevent the progression of CVB3-induced myocarditis.

## Materials and methods

### Mice

All animal experiments were performed in accordance with the National Institute of Health Guide for the Care and Use of Laboratory Animals and were approved by the Committee on the Ethics of Animal Experiments of Tongji Medical College, Huazhong Science and Technology University. six to seven week old male Balb/c mice were purchased from SPF Biotechnology Co., Ltd. (Beijing). Mice were randomly divided into four groups: the sham group (only administrated with vehicle), a group treated with the sEH inhibitor 1-(1-propanoylpiperidin-4-yl)-3-[4-(trifluoromethoxy) phenyl]urea (TPPU) (MedChemExpress LLC, 3 mg/kg per day), a group treated with CVB3 (2×10^6^ p.f.u. per g body weight), a group that received both CVB3 and TPPU (3 mg/kg per day). For infection, mice were administered (i.p.) CVB3 on day 1 to generate acute viral myocarditis and mice receiving TPPU or CVB3 and TPPU were treated with TPPU (3 mg/kg per day, dissolved in saline, i.g.) for 7 days before CVB3 infection. The sham group was administered with saline (i.p.).

Ephx2^fl/fl^ mice (NO. T009261) were purchased from GemPharmatech (Nanjing, China) and the Myh6-cre/Esr1 mice were acquired from Cyagen (Suzhou, China). The Ephx2 flox line was bred with Myh6-cre/Esr1 line to generate cardiomyocyte-specific sEH knock out (sEH^iΔMyh6^) model. Myh6-Ephx2^+/+^ littermates were used as control group. Tamoxifen (MedChemExpress LLC) dissolved in corn oil was administered intraperitoneally (i.p., 20 mg/kg per day x 5 days) to induce cre activity. Animals were treated with PBS or infected with CVB3 1 week after the final dose of tamoxifen. Exclusively male animals were studied.

### Viral infection

For viral replication assays, primary mouse neonatal cells or AC16 cells (2×10^6^) were infected with CVB3 at multiplicity of infection of 1 (MOI=1) for 1 hour, then the supernatants were removed and cells washed using PBS followed by culture in full medium for 12 or 24 hours. Viral replication was analyzed by qPCR. For mouse infection, six to seven week old male Balb/c mice were injected with CVB3 (2×10^6^ p.f.u. per g body weight). Myh6-sEH^−/−^ and Myh6-sEH^+/+^ mice were infected with CVB3 at 4×10^6^ p.f.u. per g body weight. At the end of experiments, hearts were collected for qPCR and Western blot analyses.

### Liquid chromatography-tandem mass spectrometry (LC-MS/MS)

Arachidonic acid metabolites were quantified using targeted liquid chromatography tandem mass spectrometry system (UPLC-MS/MS system, AB Sciex Triple Quad 6500+, Singapore) as described [32]. Blood samples were collected from all patients on admission and 200 μl plasma or 200 mg homogenized murine heart tissue were centrifuged (12000g, 4°C, 15minutes). Then the supernatant was extracted by ethyl acetate, the upper organic phase evaporated and the pellet was dissolved in 100 μl methanol (100%). Samples were stored at −80°C until analysis.

### Echocardiography

Cardiac function was evaluated 7 days after CVB3 infection with or without TPPU treatment by transthoracic echocardiography, using a Vevo 1100 Imaging System (Visual Sonics, Toronto, Canada), equipped with a 30 MHz linear-array transducer. M-mode images were collected in short-axis. All measurements were carried out in a blinded fashion.

### RNA extraction and real-time PCR

Total RNA was extracted from cells or mouse heart tissue using the TRIzol reagent (Invitrogen). Total RNA (500 ng) was reverse-transcribed into cDNA, and then subjected to quantative real-time PCR using SYBR Premix (Yeasen Biotechnology (Shanghai) Co., Ltd.) following the manufacturer’s instructions. The house keeping gene *Gapdh* was used for normalization. Gene expression was calculated with the 2^−ΔΔCt^ method. Primer sequences are listed in Supplemental Table 1.

### Western blotting and native PAGE

Tissues or cells were homogenized in RIPA lysis buffer (Wuhan Servicebio Tech, Wuhan, China) containing freshly added PMSF, protease inhibitors and phosphatase inhibitors (Boster Biological Tech, Wuhan, China) according to manufacturer’s instructions. The extract was centrifuged (12000 g, 15 minutes, 4°C). The protein concentration in the supernatants was determined using the BCA Protein Assay Kit (Boster Biological Tech, Wuhan, China). Proteins were separated by 10-12% SDS–PAGE. Individual proteins were detected with specific antibodies. Bands were quantified with ImageJ.

For native PAGE [33], samples in different groups were harvested with ice-cold lysis buffer (50 mM Tris-HCl, pH 8.0, 150 mM NaCl, and 1% NP-40 containing protease inhibitors). Lysis supernatant protein (12000 g, 15 minutes, 4°C) was quantified and diluted with 2x native PAGE sample buffer (125 mM Tris-HCl, pH 6.8, 30% glycerol, and 0.01% bromophenol blue). Thereafter, samples were applied to a 8% acrylamide gel without SDS at 35 mA for 50 minutes. After electrophoresis, proteins were transferred onto a nitrocellulose membrane for immunoblotting.

Primary antibodies were as follows: sEH (1:1000,10833-1-AP, Proteintech Group, Shanghai, China), IRF3 (1:1000, 4302, Cell Signaling Tech, MA, US), phospho-IRF3 (Ser396) (1:1000, 29047s, Cell Signaling Tech, MA, US), TBK1(1:1000, A3458, ABclonal Biotechnology, Wuhan, China), phospho-TBK1(Ser172) (1:1000, 5483, Cell Signaling Tech, MA, US), GSK3β(1:1000, A11731, Abclonal Biotechnology, Wuhan, China), phospho-GSK3β(Tyr216) (1:1000, ab75745, Abcam, Cambridge, UK), IFNβ (1:1000, 97450S, Cell Signaling Tech, MA, US), His-tag (1:1000, AE003, Proteintech Group, Shanghai, China), Flag-tag (1:1000, 20543-1-AP, Proteintech Group, Shanghai, China), MDA5 (1:1000, 21775-1-AP, Proteintech Group, Shanghai, China), TLR3(1:1000, 17766-1-AP, Proteintech Group, Shanghai, China), GAPDH (1:10000,10491-1-AP, Proteintech Group, Shanghai, China), Tubulin (1:2000, 11224-1-AP, Proteintech Group, Shanghai, China), β-actin (1:10000, 66009-1-lg, Proteintech Group, Shanghai, China), LaminA/C (1:1000, A0249, Abclonal Biotechnology, Wuhan, China).

### Cell culture: primary cardiomyocytes and cardiac fibroblasts

Primary neonatal mouse cardiomyocytes and cardiac fibroblasts were prepared as descried [34]. Briefly, ventricles from neonatal (1-3 days) mouse (SPF Biotechnology Co., Ltd., Beijing) were harvested and sheared into 1 mm^3^ pieces. The tissue blocks were digested with 0.075% collagenase II (Worthington, United States) at 37°C for 8-10 minites and digested cells were collected in Dulbecco’s Modified Eagle’s Medium (DMEM) containing 10% fetal bovine serum (FBS). This process was repeated several times until the heart tissue was completely digested. The harvested supernatants were pooled and centrifuged (1200 g, 5 minutes, 4°C). Cells were transferred to culture flasks and cultured for 2 hours to allow cardiac fibroblasts to adhere, while the cardiomyocytes were separated in the supernatant of the culture. Thereafter, cells were cultured in DMEM containing 10% FBS, 100 U/mL penicillin and 100 μg/mL streptomycin (37°C, 5% CO2).

HEK293 and AC16 cells were cultured in DMEM with 10% fetal bovine serum, 100 U/mL penicillin, and 100 μg/mL streptomycin. Cells were maintained at 37°C under humidified conditions, 5% CO2.

### Isolation of cardiomyocytes in adult mice

Adult cardiac myocytes were isolated using a Langendorff system as described previously [35, 36]. Briefly, mouse chest cavity was exposed and the heart was immediately removed and immersed into ice-cold Tyrode’s buffer(NaCl 125 mM, KCl 5 mM, hydroxyethyl piperazineethanesulfonic acid (HEPES) 15 mM, MgCl2 1.2 mM, and glucose 10 mM, pH 7.35–7.38 (at 25°C), saturated oxygen). Subsequently the aorta was quickly mounted on the cannula. The heart was perfused with Tyrode buffer for 5 min at 37°C. The perfusion buffer was then switched to enzymatic buffer (Tyrode buffer supplemented with 600 units /mL collagenase II), applied with constant perfusion pressure of 120 cm H2O. The perfusion was stopped when the drip rates accelerated suddenly. Then the ventricle part of the heart was cut into small tissue pieces and was pipetted gently. The digestion was neutralized using DMEM containing 10% fetal bovine serum (FBS). Then cell suspension centrifuged at 600 g for 1 minute. The CMs sediment were obtained for subsequent experiments (stored at −80°C until analyzed).

### Immunofluorescence staining

Sections (4 μm thickness) of murine hearts were permeabilized with 0.5% Triton X-100 (30 minutes, room temperature) and then blocked with 5% goat serum for 1.5 hours. Next, the sections were incubated with primary antibodies overnight at 4°C, followed by incubation with the appropriate secondary antibodies at room temperature for 1-2 hours. After washing with PBS, the sections were incubated with DAPI for nucleic acid staining within 5 minutes. Then the sections were used to take images and analysis using a ZEISS Axio Imager A2 microscope, Axiocam 506 color camera, and ZEN blue edition software.

Primary antibodies were as follows: sEH (1:200, custom made antibody generated by Proteintech Group, Shanghai, China as previously described[37]; or 1:100, sc-166961, Santa Cruz, Texas, US), cTNT (1:200, 66376-1-lg, Proteintech Group, Shanghai, China); COL1A1 (1:200, A16891, Abclonal Biotechnology, Wuhan, China), CD31 (1:200, 11265-I-AP, Proteintech Group, Shanghai, China), CD3 (1:100, ab16669, Abcam, Cambridge, UK). CD68(1:200,14-0681-80, Invitrogen, MA, US).

### Histological analysis

Mouse hearts were briefly washed in ice cold PBS and fixed in 4 % paraformaldehyde at room temperature for 2 hours before being stored at 4 °C. Hearts were embedded in paraffin and processed for histology. Hematoxylin and eosin (HE), Masson’s trichrome staining and immunohistochemistry (IHC) staining were performed according to standard procedures and analyzed for inflammation and fibrosis.

### RNA interference

Lipofectamine 2000 (Invitrogen, Waltham, MA, US) was used to transfect AC16 cells with siRNA, according to the manufacturer’s instructions. After 24 hours, cells were incubated with EETs (1 μM) or TPPU (10 μM) 1 hour before CVB3 infection (MOI=1). After 12 hours, cells were collected for subsequent detection. Human siRNA oligonucleotides targeting *MDA5, TLR3, TBK1*, and *GSK3B* (Supplemental Table 1) were from Guangzhou RiboBio Co., Ltd (Guangzhou, China).

### Co-immunoprecipitation

HEK293 cells were transfected with pc*TBK1*-His or pc*GSK3B*-Flag plasmid using Lipofectamine 2000 (Invitrogen, Waltham, MA, US) according to the manufacturer’s instructions, and treated with and without 14,15-EET (Cayman Chemical, Ann Arbor, MI, US) at 1 μM. After 48 hours, transfected HEK293 cells were harvested in immunoprecipitation (IP) buffer containing 20 mM Tris (pH7.5), 150 mM NaCl, 1% Triton X-100 with freshly added PMSF, protease inhibitors and phosphatase inhibitors (Boster Biological Tech, Wuhan, China). Immunoprecipitation was performed with anti-His or Flag magnetic beads (Bimake, Texas, US), according to the manufacturer’s instructions. Then the antigen eluted from the beads was immunoprecipitated by SDS-PAGE.

### Luciferase reporter assay

HEK293 cells (~1×10^5^) were seeded on 24-well plates and co-transfected with either empty vector (pcDNA), *pcMDA5, pcMAVS, pcTBK1* or *pcIRF3-5D* using Lipofectamine 2000 (Invitrogen, Waltham, MA, US). A p*RL-TK* renilla luciferase reporter plasmid (0.01 μg) was added to each transfection to normalize the transfection efficiency. After 4-6 hours, the supernatants were removed and cells were culture in full medium with and without 14,15-EET (1 μM). Cells were collected after 24 hours and luciferase assays were performed using a dual-specific luciferase assay kit (Promega, WI, US) according to the manufacturer’s instructions.

### Statistical analysis

Data were analyzed using the nonparametric Student’s t-test for two group comparisons; otherwise, data were analyzed by one-factor or mixed two-factor ANOVA and multiple comparisons test. Data were presented as mean SEM unless otherwise indicated in figure legends; p<0.05 was considered statistically significant.

## Results

### Levels of sEH and eicosanoid metabolites altered in myocarditis

An LC-MS/MS-based targeted approach was used to assess myocarditis-associated changes in eicosanoids including metabolites of the cyclooxygenase (COX), lipoxygenase (LOX) and CYP pathways. Interestingly, EETs were the most altered species in samples from patients with myocarditis (Fig. 1A, Fig. S1A), and their levels were significantly decreased (Fig. 1B). This correlated with an increase in the DHET/EET ratio, which is an indication of elevated sEH activity (Fig. 1C). Similar findings i.e. an increased DHET/EET ratio were made in the mouse model of CVB3-induced myocarditis (Fig. 1D-F, Fig. S1B). Indeed, CVB3 infection induced a significant increase in sEH expression that was evident as early as 12 hours after infection (Fig. S1C), clearly increased after 3 days and maintained for at least 7 days (Fig. 1G). sEH expression was largely concentrated in cardiomyocytes and fibroblasts (Fig. 1H-I, Fig.S1D).

**Fig. 1.**
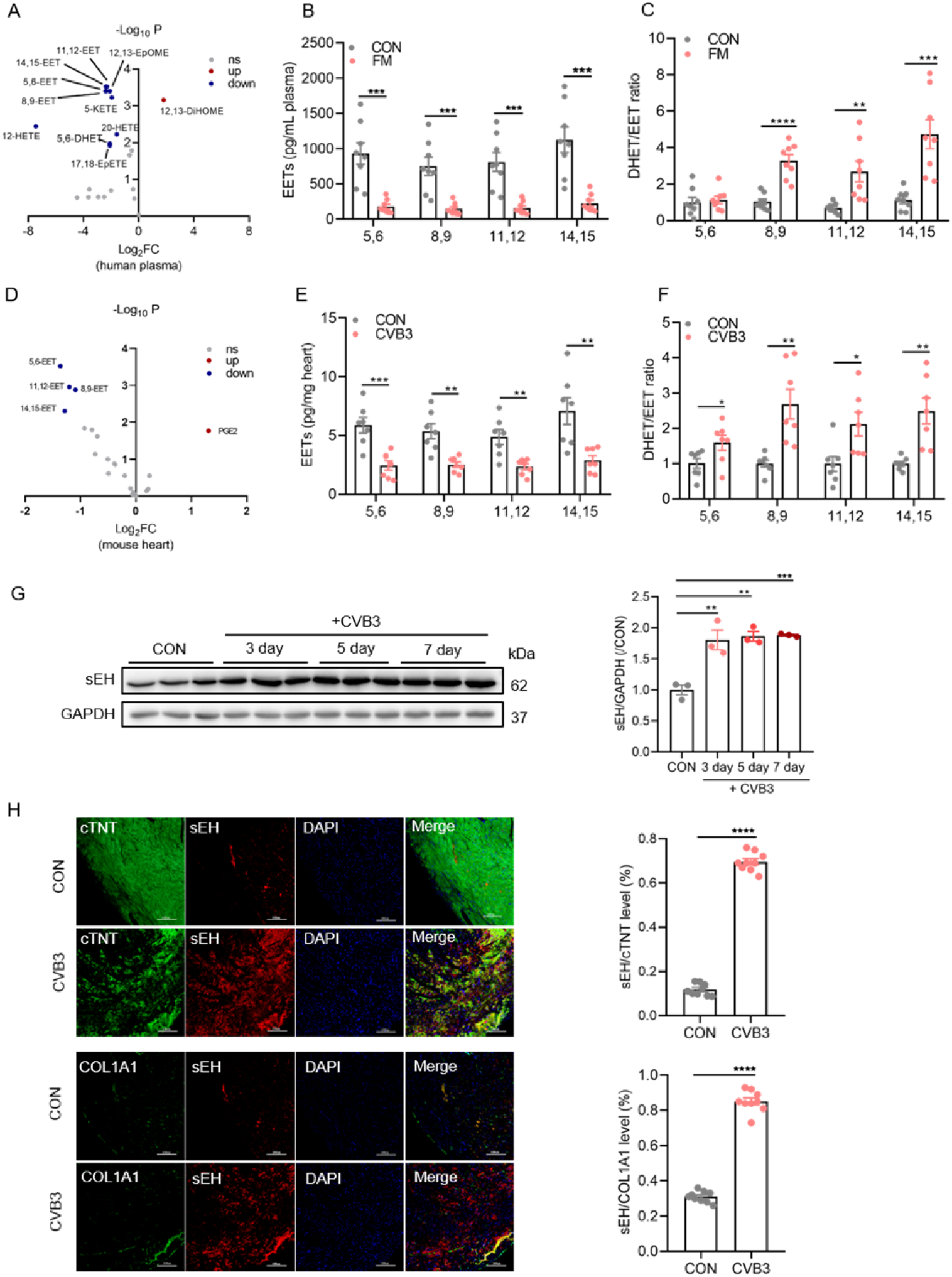
Levels of sEH and eicosanoid metabolites altered in myocarditis. (**A**) Plasma eicosanoid metabolites differentially affected in healthy subjects and patients with fulminant myocarditis (FM) (n=8). (**B-C**) Plasma levels of EETs (B), and the plasma DHET/EET ratio (C), in samples from FM patients and health subjects (n=8). (**D**) Eicosanoid metabolites in hearts from untreated and CVB3-invfected mouse hearts (n=7). (**E-F**) EET levels (E), and the cardiac DHET/EET ratio in healthy and CVB3-treated mice (n=7). (**G**) Time course of sEH expression in mouse heart for up to 7 days after CVB3-administration. GAPDH was used as the loading control (n=3). (**H-I**) Representative images showing the expression of sEH together with cTNT (H) or COL1A1 (I) in sections of mouse heart; bars =50 μm. *p < 0.05; **p < 0.01; ***p < 0.001; ****p < 0.0001.

### sEH inhibition prevents cardiac dysfunction of CVB3-induced myocarditis by promoting antiviral responses

To determine whether EETs could prevent the development of viral myocarditis, Balb/c mice were treated with CVB3 (day 0) and immediately given either PBS or the sEH inhibitor; TPPU, for up to 7 days (Fig. 2A). Compared to non-infected mice, CVB3-treated mice demonstrated a marked and continuous decrease in body weight that was less pronounced in mice given TPPU (Fig. 2B). More importantly, TPPU largely prevented the deterioration in cardiac function induced by viral infection (Fig. 2C-F), and prevented the upregulation of brain natriuretic factor (BNP), which is a marker of heart failure marker (Fig. 2G). These results demonstrated that sEH inhibition protects against the development of CVB3-induced myocarditis.

**Fig. 2.**
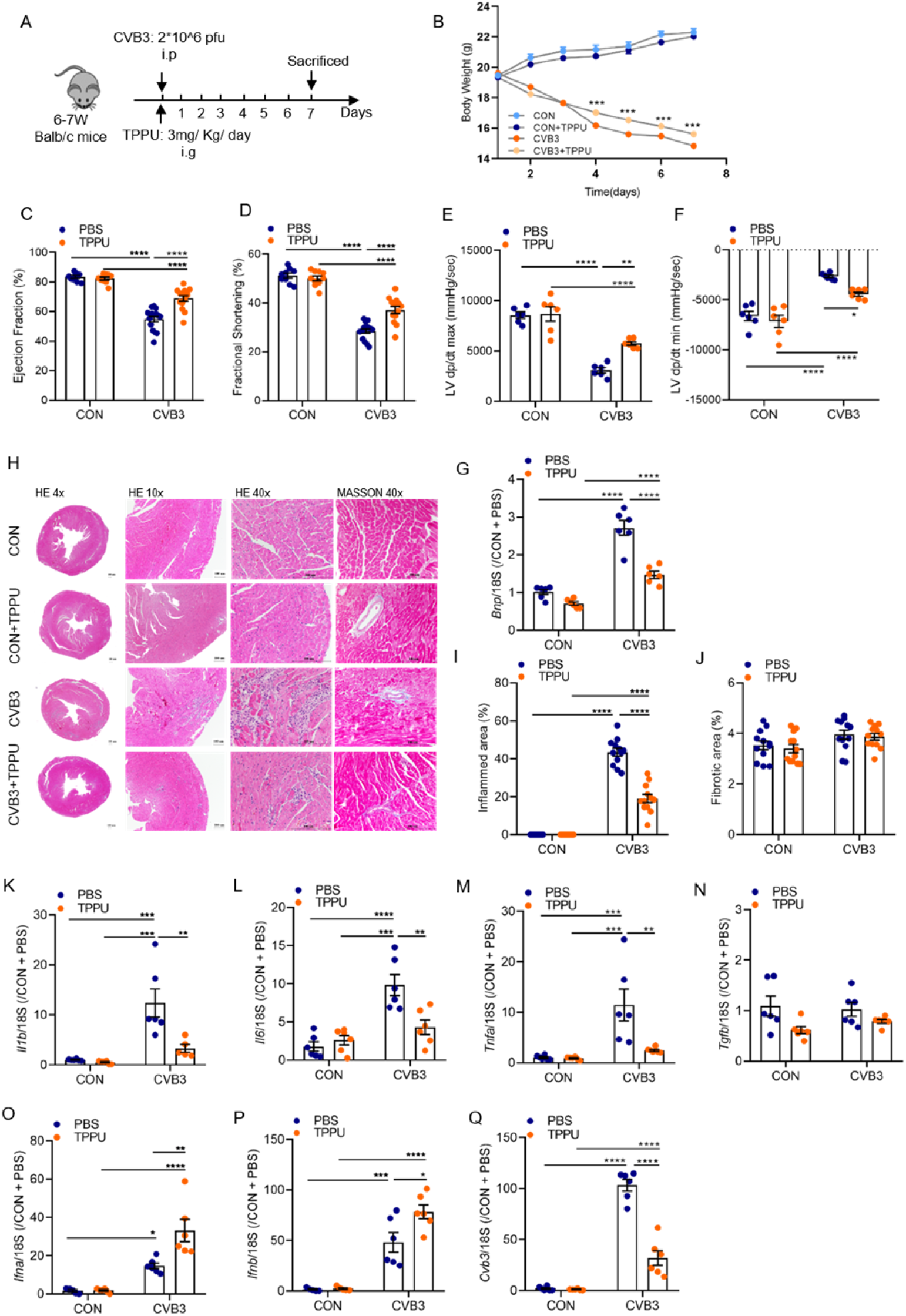
sEH inhibition prevents cardiac dysfunction associated with CVB3-induced myocarditis. (**A**) Protocol for TPPU treatment and CVB3 infection. (**B**) Body weight curves for Balb/c mice with different treatments from day 1 to day 7 (n=10-20). (**C-F**) Cardiac function in Balb/c mice 7 days after CVB3 infection detected by echocardiographic measurement and cardiac catheterization procedure (n=10-15). (**G**) Relative expression of *Bnp* (RT-qPCR). The housekeeping gene *Gapdh* was used for normalization (n=6). (**H**) Representative images showing HE and Masson’s trichrome staining of heart sections in four groups; bars =100 μm. (**I-J**) Quantification of inflamed and fibrotic areas from the images shown in H. (**K-N**) Relative expression of the pro-inflammatory mediators *Il1b, Il6, Tnfa* and *Tgfb* relative to that of the housekeeping gene *Gapdh* (n=6). (**O-P**) Relative expression of the type I IFNs; *Ifna* and *Ifnb* relative to *Gapdh* (n=6). (**Q**) Relative mRNA expression of CVB3 titers relative to *Gapdh* (n=6). *p < 0.05; **p < 0.01; ***p < 0.001; ****p < 0.0001.

As the inflammation induced by viral infection is the cause of myocardial dysfunction, we further focused on the inflammatory response and, in particular, on pathways determining host defense against viruses. Histological analysis of heart sections (H&E and IHC staining) revealed the expected increase in infiltrating inflammatory cells in mice with myocarditis that was clearly attenuated by treating mice that received TPPU (Fig. 2H-I, Fig. S2A). There was no evidence of cardiac fibrosis (MASSON staining) in the acute phase of CVB3-induced myocarditis (Fig. 2H-J). There was, however, a clear increase in mRNA levels encoding interleukin (IL)-1β, IL-6 and tumor necrosis factor (TNF)-α in hearts from CVB3-treated mice, that were again attenuated in animals that received the sEH inhibitor (Fig. 2K-N). Given the protective effects of TPPU, we also determined the expression of type I IFNs involved in the anti-viral response [21, 38]. Consistent with the effects on myocarditis, mice treated with TPPU and CVB3 expressed much higher cardiac levels of IFN-α and IFN-β than CVB3 treated mice that received PBS (Fig. 2O-P). There was no increase in IFN levels in TPPU-treated mice without the induction of viral myocarditis. Given that our data implied that sEH inhibition was able to prevent the induction of viral myocarditis, we determined whether or not sEH inhibition affected viral replication. Indeed, TPPU-treatment reduced viral titers in CVB3-infected mice (Fig. 2Q), implying that the increased production of IFNs in sEH inhibitor-treated mice prevented viral replication.

### 14,15-EET positively regulates type I IFN production

To further investigate the molecular pathways linked to sEH activity and the anti-viral response, we studied neonatal mouse cardiomyocytes and cardiac fibroblasts and infected them with CVB3 at a multiplicity of infection (MOI) of 1 for 12 hours. As in the *in vitro* model, sEH protein expression was increased by CVB3 infection in cardiomyocytes and cardiac fibroblasts (Fig. 3A, Fig. S2A), and sEH inhibition attenuated viral replication after CVB3 infection (Fig. 3B). One of the EETs most markedly affected by CVB3 was 14,15-EET, and the 14,15-EET antagonist 14,15-epoxyeicosa-5(Z)-enoic acid EEZE (1 μM), attenuated the protective effect of TPPU. Moreover, the addition of authentic 14,15-EET decreased viral replication in an EEZE-sensitive manner. Similar effects of sEH inhibition and 14,15-EET were also detected in cardiac fibroblasts (Fig. S3B-E). Next, we examined IFN expression in the different groups. As in the *in vitro* situation IFN levels were barely detectable in uninfected cardiomyocytes and pronounced in CVB3-treated cells. sEH inhibition increased the expression of IFN-α (Fig. 3C) and IFN-β (Fig. 3D) in an EEZE-sensitive manner, an effect that was more marked in cells treated with 14,15-EET (1 μM). The effects on IFN-β were more pronounced and also confirmed at the protein level (Fig. 3E). Also, the expression of the IFN-β-regulated genes i.e. ISG15 and CXCL10 were increased by CVB3 infection in a manner that was sensitive to sEH inhibition and 14,15-EET (Fig. 3F-G). Again, the effects of sEH inhibition and 14,15-EET were sensitive to the EET antagonist. The actions of IFN-α and IFN-β depend on the activation of their receptor and blockade of the IFN-α and β receptor subunit 1 (IFNAR) antagonized the antiviral properties of sEH inhibition and 14,15-EET (Fig. 3H). Thus, the anti-viral actions linked to sEH inhibition seem to be linked to 14,15-EET-dependent effects on IFN expression and IFNAR activation.

**Fig. 3.**
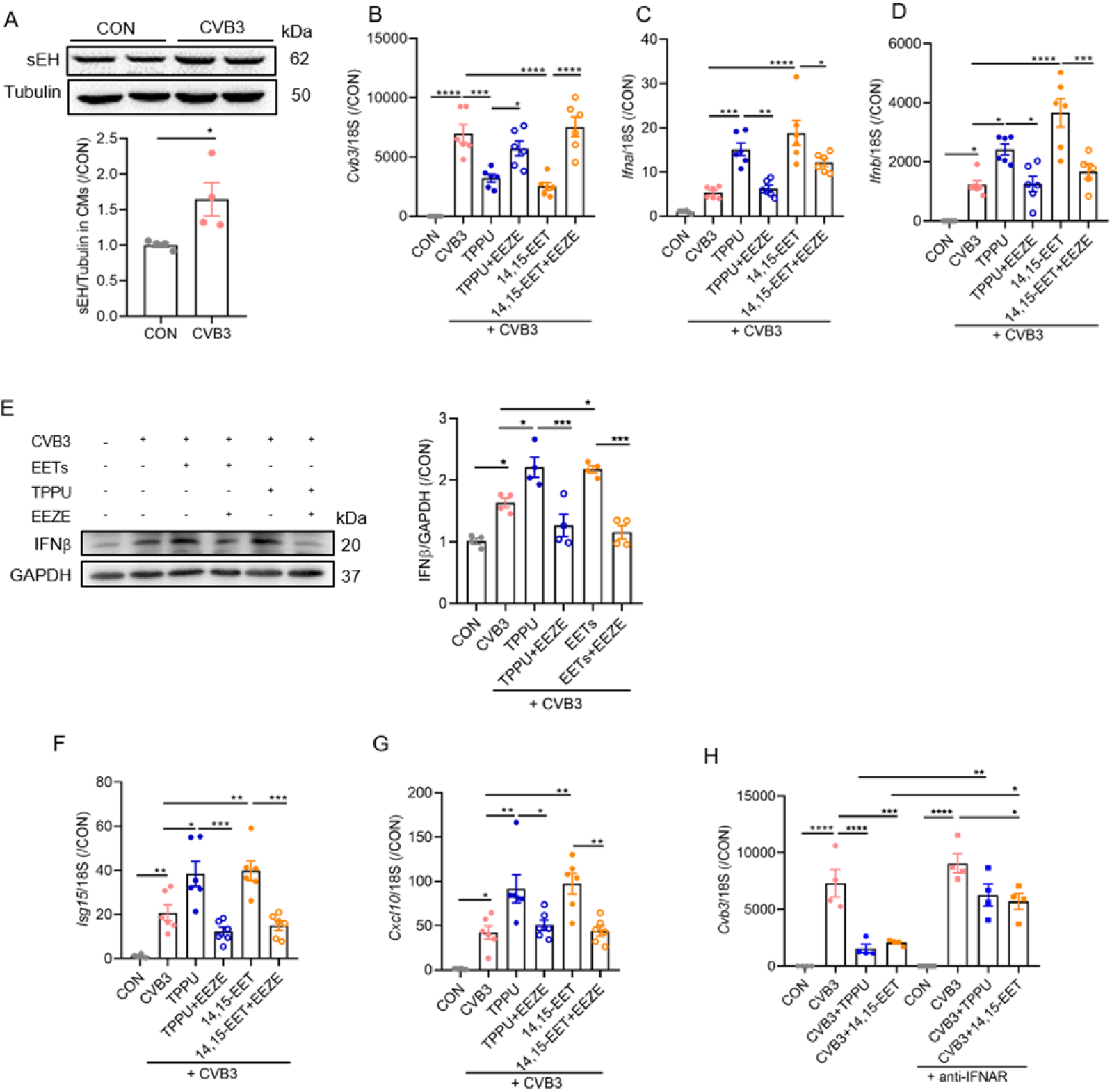
EETs/sEH inhibition positively regulate type I interferons production in vitro. (**A**) sEH expression relative to tubulin in primary cardiomyocytes from mice treated with PBS (CON) or infected with CVB3 (n=4). (**B**) Expression of CVB3 titers in cells treated with solvent (CON), TPPU (10 μM) or 14,15-EET (1 μM) alone or in combination with EEZE (1 μM, 1 hour pretreatment), 1 hour prior to CVB3 infection (n=6). (**C-D**) Expression of *Ifna* and *Ifnb* relative to *Gapdh* in animals treated as in B (n=6). (**E**) IFNβ expression in cardiomyocytes treated with solvent or infected with CBV3 in the absence and presence of TPPU (10 μM), 14,15-EET (1 μM) or EEZE (1 μM) as in B (n=4). (**F-G**) Expression of *Isg15* (F), and *Cxcl10* (G) in primary cardiomyocytes from the different groups (n=6). (**H**) Expression of CVB3 titers in cardiomyocytes treated with 14,15-EET (1 μM) or a type I interferon receptor (IFNAR) blocking antibody (10 μg/mL) 2 hours before CVB3 infection (n=4). *p < 0.05; **p < 0.01; ***p < 0.001; ****p < 0.0001.

### EETs enhanced type I interferon signaling upon CVB3 infection by targeting TBK1

Since the transcription factor; interferon regulatory factor 3 (IRF3), plays an essential role in the induction of type I IFNs in response to viral infections [39, 40], we explored a possible link between 14,15-EET and IRF3 activation. As expected, CVB3 infection resulted in the phosphorylation of IRF3 on Ser396 (Fig. 4A), an increase in the formation of active IRF3 dimers (Fig. 4B) and its nuclear translocation (Fig. 4C-D). Both the sEH inhibitor; TPPU and 14,15-EET potentiated these effects and induced a marked further increase in phosphorylation and dimer levels. The actions of 14,15-EET were also paralleled by a second EET regioisomer i.e., 11,12-EET, the levels of which were also decreased in hearts from mice with viral myocarditis (see Fig. 1A).

**Fig. 4.**
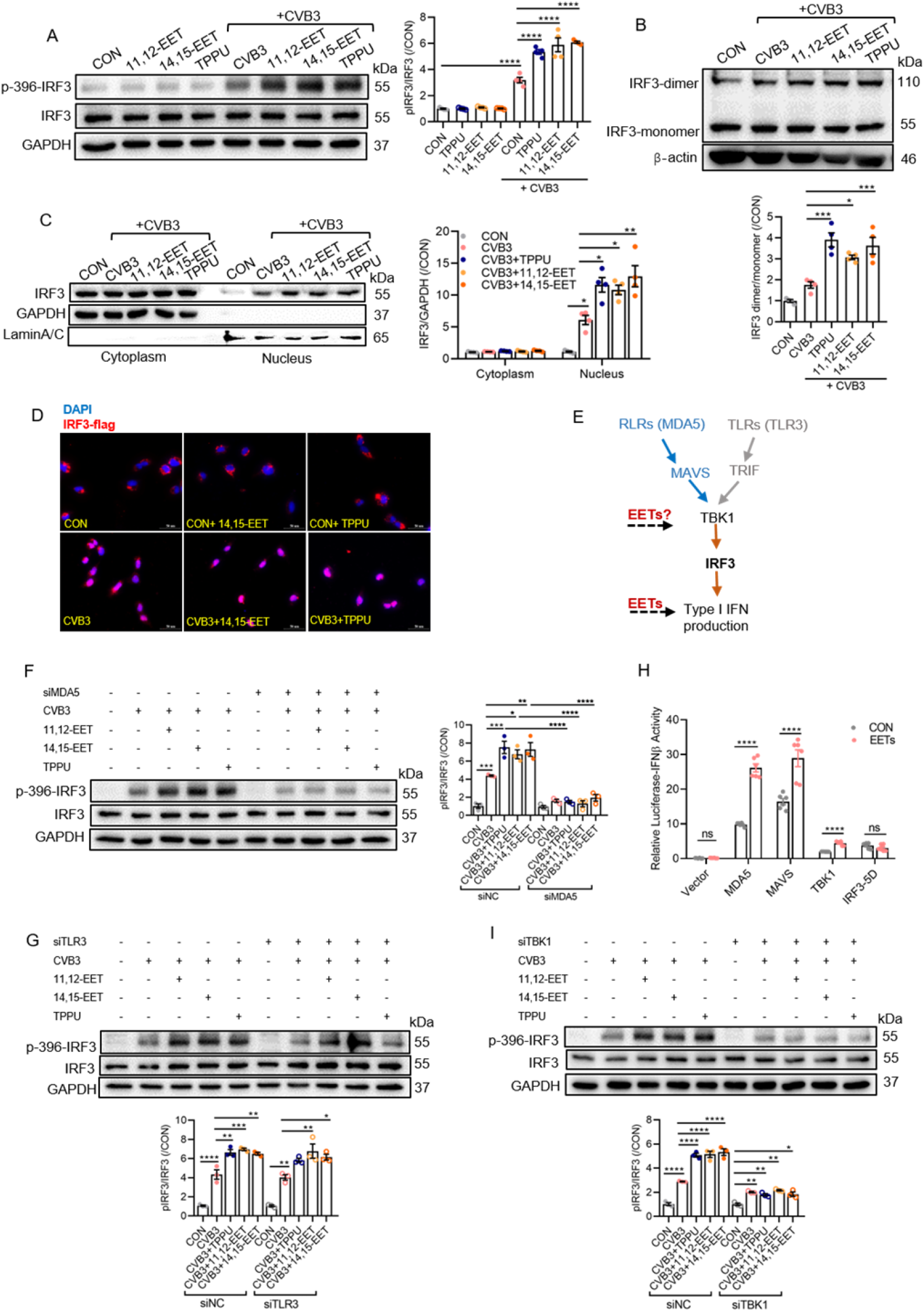
EETs enhance type I IFN signaling after CVB3 infection by targeting TBK1. (**A**) Phosphorylation of IRF3 (on Ser396) in uninfected (CON) or CVB3-infected AC16 cells treated with 11,12-EET, 14,15-EET or TPPU (n=4). (**B**) Dimerization of IRF3 in uninfected (CON) or CVB3-infected AC16 cells treated with 11,12-EET, 14,15-EET or TPPU (n=4). (**C**) Nuclear translocation of IRF3 in uninfected (CON) or CVB3-infected AC16 cells treated with 11,12-EET, 14,15-EET or TPPU (n=4). GAPDH and LaminA/C were used as cytoplasm and nuclear protein loading controls. (**D**) Immunofluorescence images showing the impact of 14,15-EET and TPPU on the subcellular localization of IRF3 in uninfected (CON) or CVB3-infected AC16 cells; bars =50 μm. (**E**) Scheme showing the two main RNA sensor-induced IRF3 activation pathways. (**F-G**) AC16 cells that were uninfected (CON) or infected with CVB3 were treated with 11,12-EET, 14,15-EET or TPPU (n=3). Shown are changes in IRF3 phosphorylation following the siRNA-mediated down regulation of MDA5 (F), or TLR3 (G). (**H**) Activity of a IFNβ-luciferase reporter construct in HEK293 cells expressing either a control vector (vector), *pcMDA5, pcMAVS, pcTBK1*, or *pcIRF3-5D* and treated with solvent (CON) or 14,15-EET (1 μM) for 24 hours. (**I**) Impact of the siRNA-mediated downregulation of TBK1 on the phosphorylation of IRF3 (on Ser396) in uninfected (CON) or CVB3-infected AC16 cells treated with 11,12-EET, 14,15-EET or TPPU (n=3). *p < 0.05; **p < 0.01; ***p < 0.001; ****p < 0.0001.

Host recognition of CVB3 is thought to be determined by melanoma differentiation-associated protein 5 (MDA5) and Toll-like receptor 3 (TLRT3) [41, 42], which activate IRF3 via Tank binding kinase 1 (TBK1) [43, 44] (Fig. 4E). The downregulation (siRNA-mediated) of MDA5 (Fig. S. 4A) effectively reduced the ability of CVB3 to elicit the phosphorylation of IRF3, as well as the ability of the sEH inhibitor, 14,15-EET and 11,12-EET to potentiate the response (Fig. 4F). The downregulation of TLR3 (Fig. S4B) was largely without consequence on the viral or EET-induced phosphorylation of IRF3 (Fig. 4G). Next, the dependence of EETs on the expression of MDA5, MAVS and TBK1 was determined in a series of IFN-β reporter-luciferase activity assays. In HEK293 cells there was no detectable activity of the IFN-β reporter but its activity was increased following the overexpression of MDA5 and its downstream target mitochondrial antiviral signaling protein (MAVS) (Fig. 4H). Similarly, the overexpression of TBK1 and a constitutively active mutant of IRF3 i.e. IRF3-5D, also increased luciferase activity. 14,15-EET was able to significantly potentiate luciferase activity in cells expressing MDA5, MAVS and TBK1 but not in cells overexpressing the IRF3-5D mutant. Silencing (siRNA-mediated) TBK1 (Fig. S4C), on the other hand, reduced IRF3 phosphorylation in response to CVB3 and abrogated the effects of the sEH inhibitor and EETs (Fig. 4 I). Overall, these findings suggest that EETs were able to promote type I IFN signaling by targeting TBK1.

### EETs positively regulated TBK1 activation via enhancing the interaction between GSK3β and TBK1

Next, we assessed how 14,15-EET and 11,12-EET were able to increase the activation of TBK1 or other signaling proteins within the type I IFN pathway *in vitro*. Neither CVB3 infection, not the treatment with TPPU, 14,15-EET or 11,12-EET had any effect on TBK1 expression (Fig. 5A), indicating that other regulatory mechanisms were involved. Because the activity of TBK1 can be regulated by multiple post-translational modifications such as ubiquitination and phosphorylation [45–48], non-infected and CVB3-infected AC16 cells were treated with 11,12-EET and 14,15-EET (1 μM) and changes in the phosphoproteome were assessed using LC-MS/MS. However, no changes in TBK1 phosphorylation were detected. Next, we immunoprecipitated TBK1 from HEK293 cells to identify co-precipitated proteins by MS. Comparative analysis of TBK1 interactome and phosphor-proteome data revealed shared and disparate proteins (Fig. 5B). A protein-protein interaction network analysis between TBK1 and the shared proteins was then performed. This step identified GSK3β as a TBK1-interacting protein (Fig. 5C, Fig. S5B-E), that was phosphorylated on Tyr216 by both 11,12- and 14,15-EET (Fig. 5D).

**Fig. 5.**
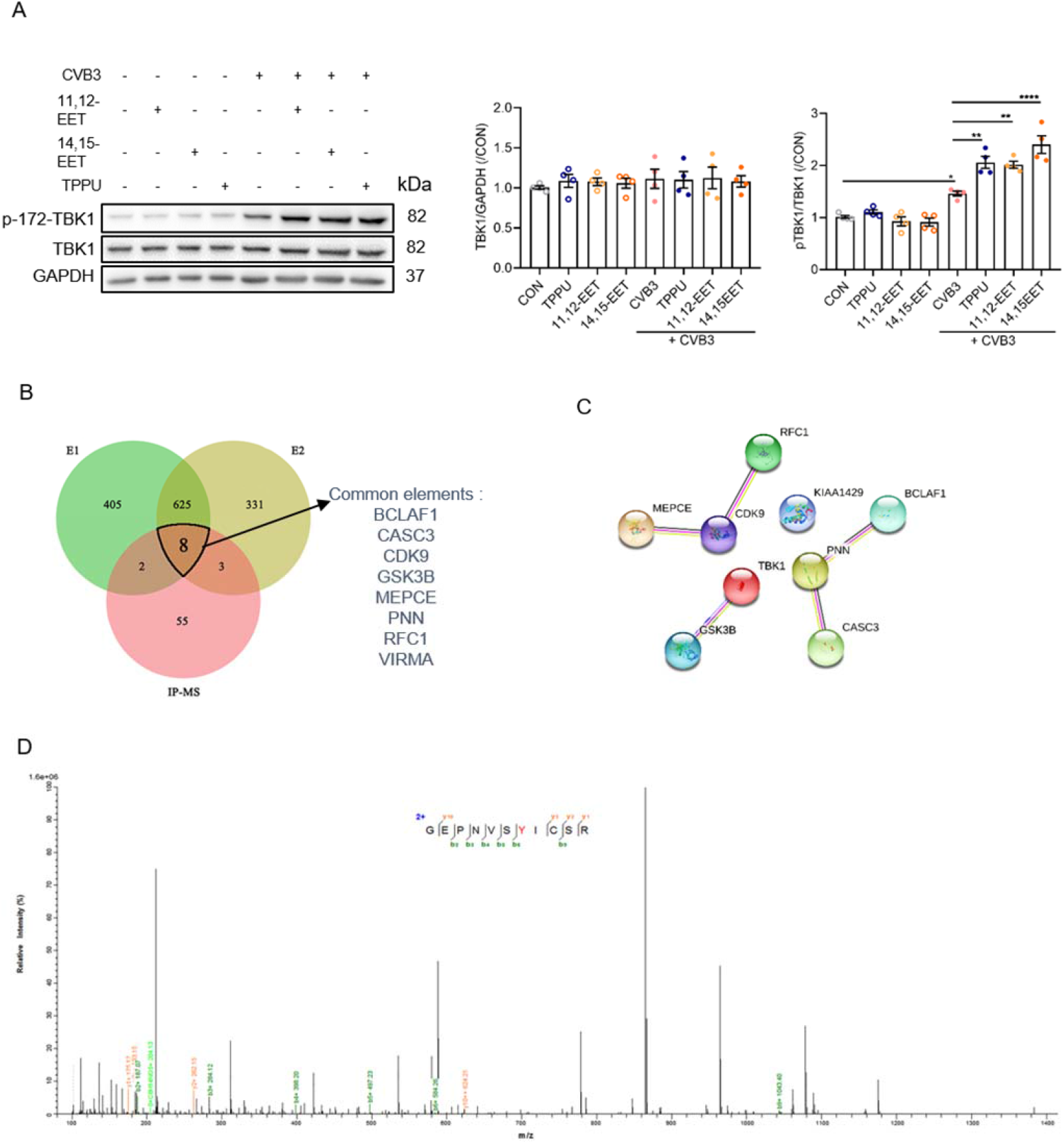
Activation of TBK1 mediated by EETs. (**A**) TBK1 phosphorylation and expression in uninfected and CBV3-infected AC16 cells treated with 11,12-EET, 14,15-EET or TPPU (n=4). (**B**) Combined analysis of the TBK1 interactome (IP-MS) and phosphoproteome from uninfected and CVB3 infected AC16 cells treated with 11,12-EET (E1) or 14,15-EET (E2) (n=3). (**C**) STRING analysis reveal potential protein interaction network among TBK1 and targets from (B). (**D**) Detection of phosphorylated residues in GSK3β immunoprecipiteated from AC16 cells treated with 14,15-EET (n=3). *p < 0.05; **p < 0.01; ****p < 0.0001.

GSK3β was essential for the EETs to increase IFN signaling (IRF3 phosphorylation) in CVB3-infected AC16 cells as its silencing (Fig. S5A) prevented the response (Fig. 6A). Importantly, IRF3 phosphorylation stimulated by CVB3 infection alone was unaffected thus suggesting that the actions of the EETs were dependent on the expression of the kinase. It is not entirely clear how EETs are able to elicit their effects but a considerable amount of evidence indicates that a specific EET receptor exists on the cell membrane [49–51]. Indeed, the ability of a series of EET antagonists to abrogate EET-induced signaling events is strong evidence for the existence of such a receptor. In AC16 cells, CVB3 infection effectively increased the tyrosine phosphorylation of GSK3β. This response was potentiated by 11,12- and 14,15-EET and antagonized by EEZE indicating the involvement of an extracellular event (Fig. 6B). In addition to the phosphorylation of GSK3β, 14,15-EET enhanced the association of GSK3β with TBK1 under CVB3 infection (Fig. 6C). Therefore, we generated non-phosphorylatable and phosphomimetic GSK3β mutants by replacing Tyr216 with aspartate (Y216D) or phenylalanine (Y216F), respectively. Interestingly, the Y216D mutant resulted in the marked phosphorylation of TBK1 only in virally infected cells (Fig. 6D). Consistent with these observations, 14,15-EET increased TBK1 phosphorylation in CVB3-infected cell expressing the wild-type GSK3β but not in cells expressing the non-phosphorylatable GSK3β mutant (Fig. 6E-F). Thus, in the course of CVB3 infection, EETs are able to positively regulate TBK1 activity by eliciting the phosphorylation of GSK3β phosphorylation to enhance its interaction with TBK1.

**Fig. 6.**
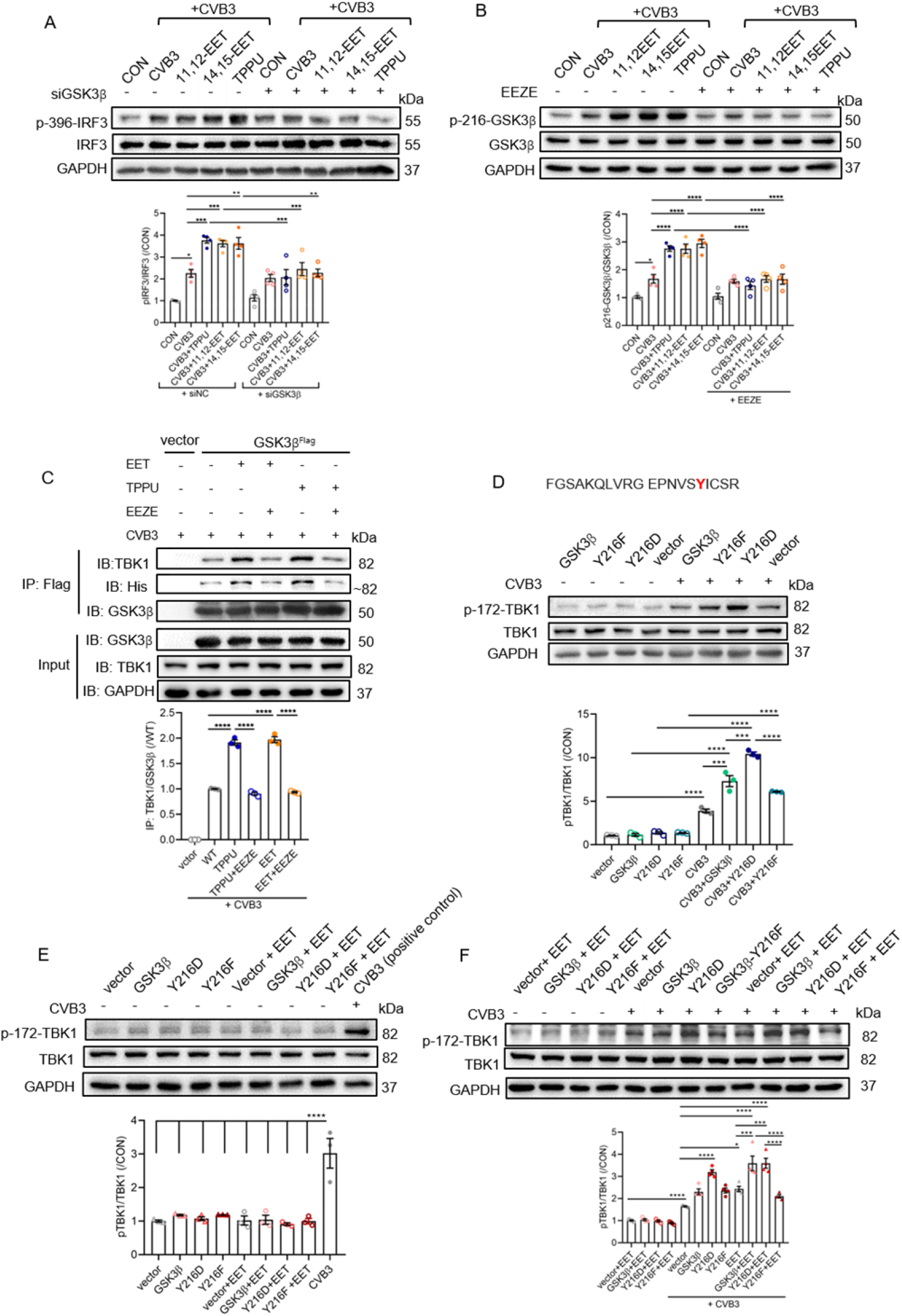
EETs positively regulated TBK1 activation via enhancing the interaction between GSK3β and TBK1. (**A**) Impact of the siRNA-mediated downregulationof GSK3β on the phosphorylation of IRF3 (on Ser396) in uninfected (CON) or CVB3-infected AC16 cells treated with 11,12-EET, 14,15-EET or TPPU (n=4). (**B**) Impact of the EET antagonist; EEZE (1 μM) on the phosphorylation of GSK3β in AC16 cells treated with 11,12-EET, 14,15-EET or TPPU (n=4). (**C**) Co-immunoprecipitation of TBK1 with a GSK3β-Flag fusion protein from HEK293 cells treated with 14,15-EET (1 μM) or TPPU (10 μM) in the absence or presence of EEZE (n=3). (**D**) Impact of the wild-type GSK3β and the Y216F and Y216D GSK3β mutants on the phosphorylation of TBK1 in uninfected and CVB3-infected HEK293 cells (n=3). (**E-F**) Impact of GSK3β mutation alone and in combination with 14,15-EET on the phosphorylation of TBK1 in uninfected and CVB3-infected HEK293 cells (n=3-4). *p < 0.05; **p < 0.01; ***p < 0.001; ****p < 0.0001.

To further investigate the cardioprotective effect of EETs on cardiomyocytes in vivo, we generated cardiomyocyte-specific sEH knockout (sEH^iΔMyh6^) mice by crossing Myh6-CreEsr1 mice with Ephx2^fl/fl^ mice and ablated Ephx2 expression by treating the mice with tamoxifen (Fig. 7A, Fig. S6A-F). One week after the knockdown of cardiomyocyte sEH expression, animals were infected with CVB3. Similar to the responses observed in wild-type mice given the sEH inhibitor, the cardiomyocyte specific deletion of the enzyme attenuated the virus-induced decrease in cardiac function (Fig. 7B-C). Subsequent, histological analysis revealed that sEH deletion significantly reduced the inflammatory reaction caused by CVB3 (Fig. 7D-E, Fig. S6G-H). To link the latter observations with the data generated in the cell line, adult mouse cardiomyocytes were isolated from mice 7 days after CVB3 infection. This revealed increased phosphorylation of TBK1 and GSK3β in cardiomyocytes from sEH^iΔMyh6^ mice (Fig. 7F-H).

**Fig. 7.**
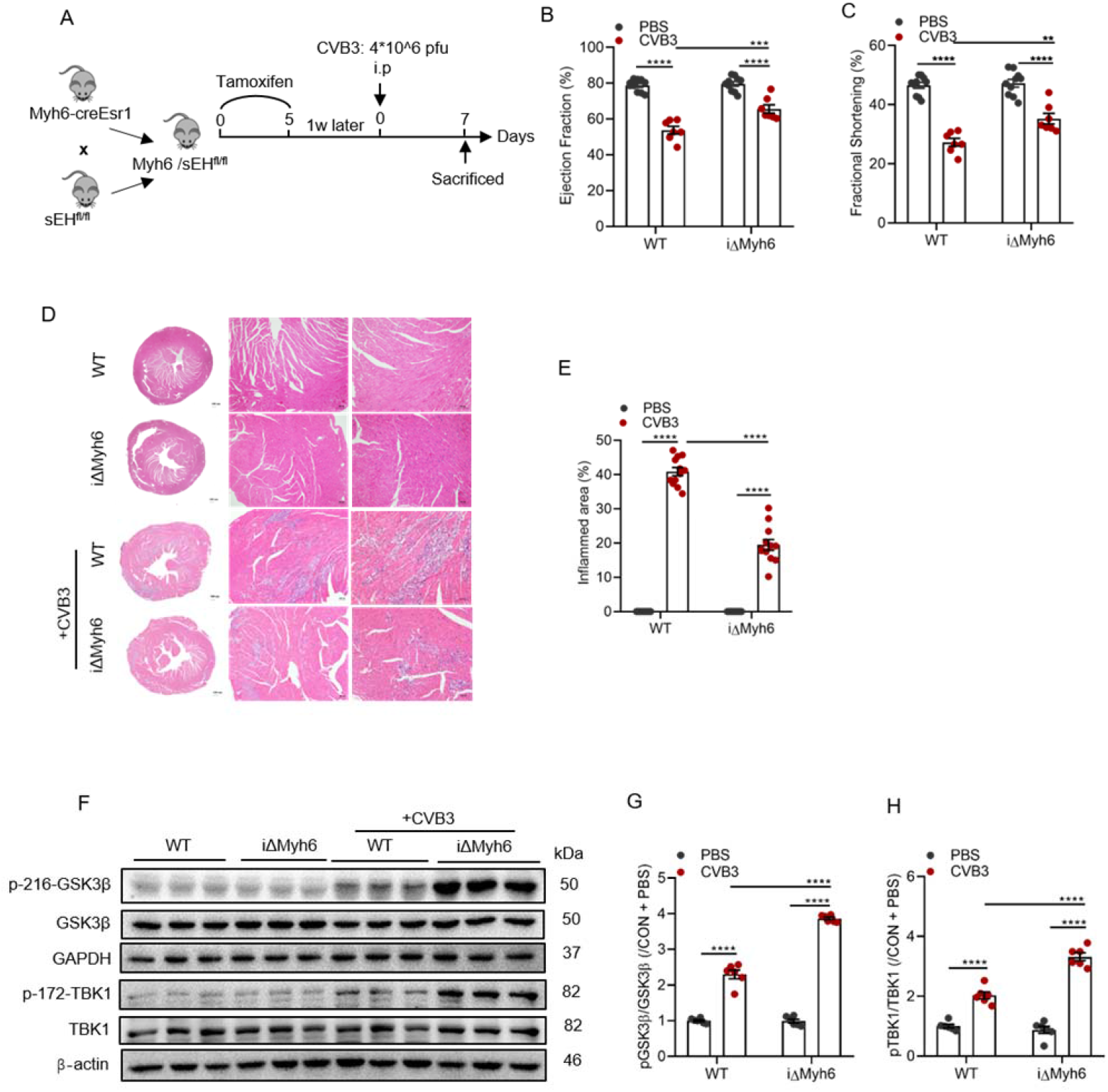
Impact of cardiomyocyte-specific sEH deletion on cardiac function following CVB3 infection in vivo. (**A**) Scheme showing the tamoxifen application protocol prior to CVB3 infection. (**B-C**) Cardiac function (echocardiography) in wild-type (WT) and sEH^iΔMyh6^ (iΔMyh6) littermates 7 days after being given PBS or infected with CVB3. Shown are ejection fraction (B) and fractional shortening (C); n=7-10. (**D-E**) Infiltrating cells in hearts from wild-type (WT) and sEH^iΔMyh6^ (iΔMyh6) littermates 7 days after being given PBS or infected with CVB3 (n=4); bars =100 μm. (**F-H**) Phosphorylation of GSK3β and TBK1 in cardiomyocytes from wild-type and sEH^iΔMyh6^ (iΔMyh6) littermates 7 days after PBS or CVB3 infection (n=6). *p < 0.05; **p < 0.01; ***p < 0.001; ****p < 0.0001

## Discussion

The results of our investigation revealed that increasing EET levels; by inhibiting the sEH or by their exogenous application, stimulates type I IFN production and IFN signaling to attenuate the myocardial dysfunction induced by CVB3-infection. The first evidence that the sEH could play a role in viral myocarditis was that EET levels were decreased in plasma from patients with fulminant myocarditis compared to healthy controls. In mice infected with CVB3, it was possible to demonstrate that EET levels were also reduced in the myocardium and that this correlated with an increase in sEH expression. Importantly, treating animals with a sEH inhibitor increased EETs, decreased viral load and improved cardiac function. At the molecular level, we investigated the mechanism concerning up-regulated expression of IFNs by EETs. We found that EETs promoted the interaction of GSK3β with TBK1 to promote the activity of the kinase, thereby enhancing IRF3-mediated type I IFN signaling.

In the murine heart, sEH levels were low under control conditions but clearly upregulated after viral infection and most prominent in cardiomyocytes and fibroblasts. To investigate to what extent the cardioprotective effect of the sEH inhibition relied on sEH expression in cardiomyocytes, we generated sEH^iΔMyh6^ mice and infected them with CVB3 1 week after the tamoxifen-induced deletion of the enzyme. The cardiomyocyte specific deletion of sEH attenuated prevented the virus-induced decrease in cardiac function as well as damping the inflammatory cell infiltration and the generation of inflammatory cytokines. Following CVB3 infection, pattern-recognition receptors such as the Toll-like receptors and retinoic acid-inducible gene-I like receptors (RLRs) recognize viral RNAs to activate type I IFN signaling [15, 41, 42]. Several previous studies investigated IFN expression in response to viral infection of cardiomyocytes and cardiac fibroblasts and reported that cardiomyocytes express higher levels of IFNs under basal conditions while IFN production by cardiac fibroblasts tended to be more sensitive to viral infection [52–55]. In this study, we found that CVB3 infection increased cardiac IFN production and that sEH inhibition augmented this response. Thus, our observations not only confirmed the importance of investigating cell-type-specific innate immune response for organ protection [55, 56], but also identified a novel antiviral action of EETs in cardiomyocytes. Our results can also account for the recently reported actions of 11,12-EET on herpes simplex virus-infected macrophages[57].

Numerous links have been made between the CYP-sEH pathway and inflammation. For example, inflammation was reported to decrease the expression of CYP enzymes, which attenuated EET production. Given the reported anti-inflammatory actions of cellular EETs [27], this effect on its own would tend to promote inflammation [58]. How the EETs exert their anti-inflammatory actions remains to be fully elucidated as the initially reported EET-mediated inhibition of NF-κB [27], is context specific and increased reactive oxygen species generation associated with the activity of some CYPs could actually culminate in net NF-κB activation[59]. Other anti-inflammatory pathways targeted by EETs involve PPARα/γ and HO-1 activation [29]. Indeed, 14,15-EET was able to attenuate the LPS-induced activation of macrophages (M1 polarization) and promote their alternative (M2) activation via PPARα/γ [29]. In this study it was possible to identify a link between EETs (11,12- and 14,15-EET) and the activation of IFN signaling. While some studies implicated a critical role of Toll-like receptor signaling in IFN production in response to CVB3 infection [60, 61], we found that 14,15-EET positively regulated IRF3 activation and IFN production mainly via RLR activation. One possible reason for this discrepancy is that the involvement of Toll-like receptors in the viral response differs between cell types [62].

In our model, the phosphorylation and activation of TBK1 was downstream of RLR. Given that the activity of TBK1 is regulated by posttranslational modification i.e., ubiquitination and phosphorylation [45–48], we further investigated the regulation of TBK1 activation by EETs. While there was no evidence for a direct effect of 14,15-EET on TBK1 we found that 14,15-EET promoted the phosphorylation of GSK3β (on Y216) in CVB3 infected cells. GSK3β is of relevance as in addition to its involvement in the regulation of glycogen metabolism [63], it is also involved in IFN induction [47, 48, 64–66]. The tyrosine phosphorylation of GSK3β in EET-treated, CVB3-infected cells promoted the activation of TBK1, as well as the subsequent phosphorylation of IRF3; a step that is essential for the induction of type I IFNs [39, 40]. While the intracellular events that occurred after GSK3β activation could be elucidated, exactly how 11,12- and 14,15-EET was able to activate GSK3β remains to be determined. However, given that a so-called EET antagonist i.e. EEZE, was able to prevent the EET-induced tyrosine phosphorylation of GSK3β, it is tempting to speculate the involvement of an extracellular EET receptor[67]. Taken together our results provide convincing evidence for an antiviral action of both endogenously generated and exogenously applied EETs and imply that because of the IFN bosting propertied sEH inhibitions should be considered as a new therapeutic strategy to complement existing treatments of viral myocarditis.

**Figure 8.**
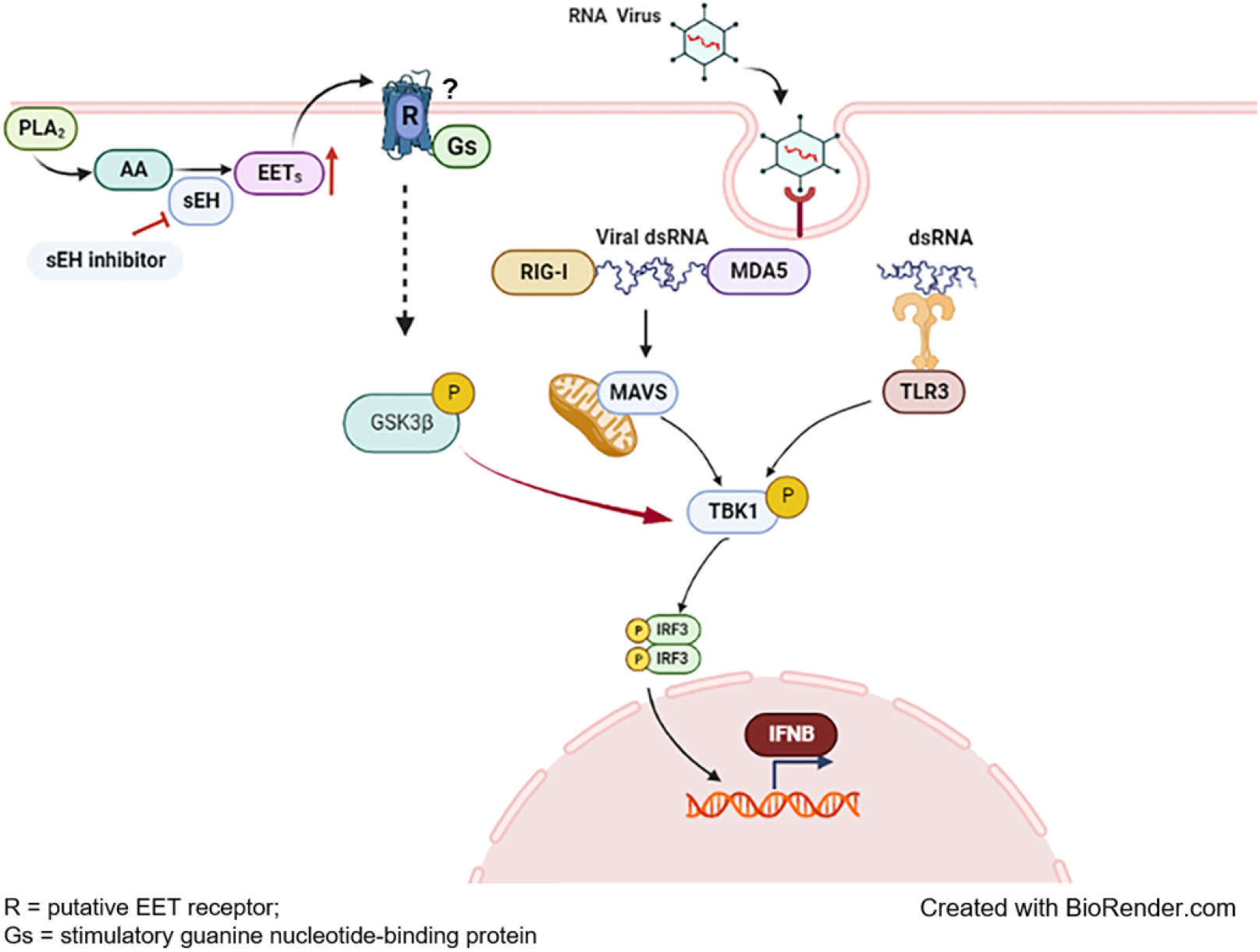

## Author contributions

Z.Z. designed and performed the majority of experiments, and drafted the paper. M.Z., C.Z. and X.G. helped to perform in vitro experiments. Z.W. performed echocardiography measurements. J.W. contributed to arachidonic acid metabolites analyses, C.C., J.H. and I.F. reviewed and revised the draft. D.W.W. designed, analyzed, and revised the manuscript. All authors read and approved the final manuscript

## Acknowledges

This study was supported in part by projects from National Nature Science Foundation of China (Nos. 81790624 and 81630010), Deutsche Forschungsgemeinschaft (SFB 1531/1 B03; Project ID: 456687919) and Top-Notch Talent Program of Hubei Province and Tongji Hospital (No. 2021YBJRC005). The funding sponsors played no role in study design, data collection and analysis, interpretation, writing of the report, and the decision to submit the paper for publication.

